# Systematically investigating the key features of the nuclease deactivated Cpf1 for tunable multiplex genetic regulation

**DOI:** 10.1101/297903

**Authors:** Chensi Miao, Huiwei Zhao, Long Qian, Chunbo Lou

**Affiliations:** CAS Key Laboratory of Microbial Physiological and Metabolic Engineering, State Key Laboratory of Microbial Resources, Institute of Microbiology Chinese Academy of Sciences, Beijing, 100101, China; College of Life Science, University of Science and Technology of China, Hefei 230027, China; Department of Biology and Center for Genomics and Systems Biology, New York University, 12 Waverly Place, New York, New York, 10003, USA.; University of Chinese Academy of Science, Beijing, 100149, China

## Abstract

With a unique crRNA processing capability, the CRISPR associated Cpf1 protein holds great potential for multiplex gene regulation. Unlike the well-studied Cas9 protein, however, conversion of Cpf1 to a transcription regulator and its related properties have not been systematically explored yet. In this study, we investigated the mutation schemes and crRNA requirements for the nuclease deactivated Cpf1 (dCpf1). By shortening the direct repeat sequence, we obtained genetically stable crRNA co-transcripts and improved gene repression with multiplex targeting. A screen of diversity-enriched PAM library was designed to investigate the PAM-dependency of gene regulation by dCpf1 from *Francisella novicida* and *Lachnospiraceae bacterium*. We found novel PAM patterns that elicited strong or medium gene repressions. Using a computational algorithm, we predicted regulatory outputs for all possible PAM sequences, which spanned a large dynamic range that could be leveraged for regulatory purposes. These newly identified features will facilitate the efficient design of CRISPR-dCpf1 based systems for tunable multiplex gene regulation.

## Introduction

Ever since the discovery of the Clustered Regularly Interspaced Short Palindromic Repeats (CRISPR) mechanism, its DNA-targeting strategy has been extensively characterized and masterfully adapted to a biotechnological tool for sequence-specific DNA manipulation that has rapidly revolutionized the fields of genome editing and DNA assembling (1-4). This simple yet elegant system consists of the Cas9 endonuclease from *Streptococcus pyogenes* and a guide RNA (gRNA) that directs Cas9 to the complementary DNA target in the presence of a protospacer adjacent motif (PAM) (2,4). The programmability, achieved through the guide sequence, has been further leveraged in variants of the system utilizing the engineered nuclease-deactivated Cas9 (dCas9) on its own or linked to diverse effector protein domains (5). These dCas9-based CRISPR toolkits have proven extremely powerful for systematic perturbation of single genes in regulatory and metabolic networks, advancing our knowledge in synthetic and systems biology at an unprecedented speed (6,7).

To push forward the CRISPR technology to the systems level, the ability to simultaneously manipulate multiple genes is highly demanded. Multiplex gene targeting, ideally through co-expressing multiple gRNAs in the same cell, enables the interrogation of much more complex interactions in genome-scale networks (8,9), as well as the combinatorial optimization of large heterologous pathways for metabolic engineering (5,8-12). Such endeavors are however hindered by current technical hurdles of multiple gRNA co-transcription – in particular, by a lack of sequence-specific RNA processing factors that do not impose significant cytotoxicity (13). This conundrum may now be solved thanks to the discovery of Cpf1, a CRISPR endonuclease of Type V-A, which displays endoribonuclease activity and was shown to process CRISPR RNA (crRNA) co-transcripts into independent mature crRNAs, in addition to its DNA cleavage activity (14-16). We thus believe in the great potential of a nuclease deactivated Cpf1 (dCpf1) as an efficient tool for multiplex gene regulation.

Although aspects of the CRISPR-Cpf1 system as DNA endonuclease has been characterized, there have been only first attempts in using CRISPR-dCpf1 as transcriptional regulators. These studies proved its applicability in bacterial, plant, and mammalian cells (17-20). To harness and streamline the system for multiplex gene regulation, three specific aspects need addressing or systematic characterization: 1. a mutational scheme that abolishes Cpf1’s nuclease activity and yet minimally affects its DNA binding and RNase activities; 2. the requirements for pre-crRNA that contains multiple direct repeat-guide sequence units for efficient crRNA processing and DNA targeting (14,21); and 3. the dependence of DNA binding strength on the PAM sequence (22-25).

In this study, we designed a negative reporter assay for transcriptional repression by the CRISPR-dCpf1 system in *Escherichia coli*. The reporter assay was used to systematically quantify the functional effects of dCpf1 mutations and crRNA variants. We evaluated the dependence of gene repression on crRNA processing, lengths of direct repeats and guide sequences, as well as the number of target sequences tandemly located within the target gene. We further investigated the PAM sequence preference for dCpf1 from *Francisella novicida* and *Lachnospiraceae bacterium* in a randomized 6nt PAM library. We found a broad range of repression activity that did not conform to the previously identified PAM preferences. Therefore, we built an interpolation algorithm to predict gene repression activity for any PAM sequence based on a much limited number of sampled weak and strong PAMs. Without assuming context dependency, the algorithm generated reliable estimates of PAM strengths, which could in principle lends great controllability and predictability to the CRISPR-dCpf1 system in synthetic biological applications.

## Materials and Methods

### Strains and Media

The *E. coli* DH5α was used as the host strain for all experiments. Luria-Bertani (LB) media (10 g/L tryptone, 5 g/L yeast extract, 10 g/L NaCl) is used as the growth media. Cells for flow cytometric fluorescence analysis were cultured in M9 media (12.8g/L Na2HPO4.7H2O, 3g/L KH2PO4, 0.5g/L NaCl, 1.67g/L NH4Cl, 1mM thiamine hydrochloride, 0.4% glucose, 0.2% casamino acids, 2mM MgSO4, 0.1mM CaCl2). Ampicillin, Kanamycin and Chloramphenicol concentrations for all experiments were 100 μg/ml, 50 μg/ml and 20 μg/ml, respectively.

### Plasmid Construction

The FnCpf1 gene were synthesized by Genscript Inc. Then it was mutated into dFnCpf1 and inserted into a vector containing a pTac-inducible promoter, an ampicillin-selectable marker, and a p15A replication origin. The crRNA plasmid backbone contained a synthetic constitutive promoter (J23119), a chloramphenicol-selectable marker, and a ColE1 replication origin. Various guide sequences were inserted by the Golden Gate method. The reporter plasmid contained *sf-gfp* as the reporter gene under the control of a synthetic constitutive promoter (J23100), a KanR-selectable marker, and a pSC101 replication origin. The crRNA sequences used in this study was summarized in Tables S3, S4 and S5.

### Flow Cytometry and Analysis

Overnight culture of *E. coli* DH5α containing test plasmids was diluted 196 times into M9 medium with corresponding antibiotics, followed by shaking at 37°C for 3 hours. Cells were then serially diluted 1000 times into M9 medium with antibiotics and appropriate concentrations of IPTG cultured at 37°C. The levels of fluorescence protein were analyzed by BD^TM^ LSR II flow cytometer (Becton Dickinson, San Jose, CA, USA) with appropriate voltage settings (FSC:440, SSC:260, FITC:480) after further dilution into PBS with 20 mg/mL Kanamycin. Each sample was collected at least 50,000 events. The mean fluorescence of each sample was calculated with Flowjo software (Treestar, Inc., San Carlos, CA, USA) and analyzed with GraphPad Prism software (GraphPad Software, La Jolla, CA, USA).

### PAM screen and Analysis

Randomized PAM library was constructed by reverse PCR and Gibson ligation, using Random_F /Random_R consisting of six randomized nucleotides as primers and plasmid R_PAM as the backbone (Figure S3). The PAM plasmid library was then transformed into competent *E.coli* DH5α harboring dFnCpf1 and crRNA plasmids. After transformation, cells were plated on LB agar supplemented with antibiotics of ampicillin, chloramphenicol and Kanamycin. After ~16 hours of growth, >10^7^ cells were collected and pooled, diluted into fresh LB medium with antibiotics, and cultured overnight (~16h). The overnight culture was diluted ~500 times into M9 medium with required antibiotics and appropriate concentrations of IPTG, followed by shaking at 37°C for 3 hours. Cultures were then diluted into PBS buffer to sort the cells with lowered fluorescence on a BD Influx Cell Sorter (Becton Dickinson, San Jose, CA, USA). From the sorted cells, random samples were collected, diluted and coated, and the remaining cells were cultured for the next round of sorting. After three rounds of sorting, colonies on the coated plates from all rounds were picked and subject to fluorescence measurements by flow cytometry and Sanger sequencing for their respective PAM sequences (Figure S4).

### PAM strength prediction and algorithm evaluation

The computation algorithm used to predict PAM strength was explained in detail in Supplementary Information. The code was written in Matlab®. Cross-validation was done by randomly selecting samples from measured mean values to generate training sets. Testing was done on measured mean values for unselected PAMs (testing sets). For original selection, samples were selected randomly from the original data set. For uniform selection, samples were selected with equal numbers from equally placed bins in the entire fluorescence range of the original data set. At each training-testing set splitting ratio, 100 independent runs were conducted. At low training set sizes, unpredicted words were removed from correlation calculations. Sequence logos in Figures 6A & S6A were generated on http://weblogo.berkeley.edu/logo.cgi.

## Results

### Single mutation dCpf1 elicits stronger gene repression than double mutation dCpf1

A previous study identified key amino acids in the RuvC-like domain of Cpf1 and proposed a double mutation scheme (D917A and E1006A) for deactivating the nuclease activity of Cpf1 from *F. novicida* (FnCpf1), in much the same way as the design of dCas9 (24). However, unlike Cas9, single mutations of either amino acid in Cpf1 was able to abolish cleavage of both DNA strands, and Cpf1 has a more complex domain structure than Cas9. We suspected that double mutations may interfere with the RNA processing and DNA binding abilities of Cpf1 and thereby affect its regulatory activity. Therefore, we constructed single mutation forms of Cpf1 nucleases from *F. novicida* (dFnCpf1) and *L. bacterium* (dLbCpf1), and tested their gene repression activities against the double mutation forms. The repression activity was tested by a negative reporter assay where a constitutively expressed *sf-gfp* gene was targeted in its promoter region by a crRNA. Upon induction of the dCpf1 variants by IPTG, reduction in fluorescence was measured as a proxy for the binding strength of the dCpf1-crRNA duplex to the DNA target (Figure 1A). Figures 1C and D show the repression activity as a function of inducer concentration for dFnCpf1 and dLbCpf1, respectively. High levels of dCpf1 led to drastic reductions in *gfp* expression; but at all concentrations, at least one of the single mutation dCpf1s out-performed the double mutation variants. At the saturating induction level, both single mutation dLbCpf1s (D832A and E925A) showed slightly but significantly higher (>2-fold) repression activity as the double mutation dLbCpf1 (D832A+E925A). For dFnCpf1, the single mutation variant D917A elicited >200-fold gene repression, followed by the double mutation variant D917A+E1006A causing strong repression as well, whereas repression by the single mutation variant E1006A was moderate, suggesting E1006A might have destabilized DNA binding but this effect was apparently compensated by the D917A mutation in the double mutation dFnCpf1 (Figure 1B-D). Antibiotic resistance borne on the *sf-gfp* plasmid was not compromised in clones carrying the single mutation dCpf1s, suggesting the enhanced repression activity was not a result of the disruption of *sf-gfp* gene sequence by residue DNase activities (data not shown). These data revealed a conserved D at position 917/832 responsible for the nuclease activity and its minimal interference with DNA binding ability. Thus, we adopted the single mutation dCpf1s *(i.e*. D917A for dFnCpf1 and D832A for dLbCpf1) in the following experiments.

**Figure 1.**
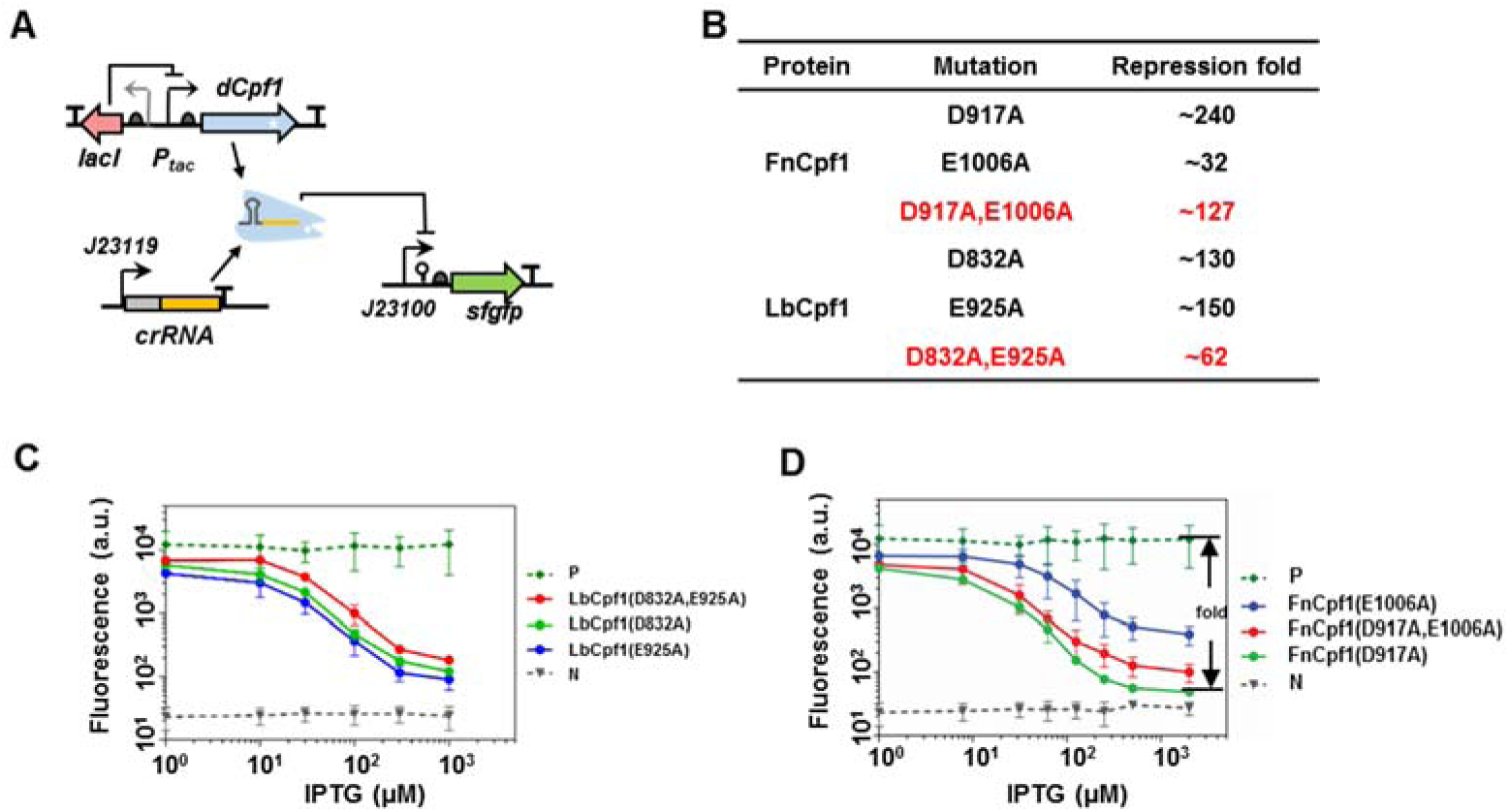
Mutation variants of dCpf1 induced differential gene repression. (A) Schematic representation of the cellular circuit for evaluating performance of the dCpf1-crRNA system. In the circuit, dCpf1 and crRNA were expressed from an inducible promoter (Ptac) and a constitutive promoter (J23119), respectively, and a reporter gene (super-folded gfp, *sf-gfp)* is repressed by the dCpf1-crRNA complex at promoter and transcribed regions. (B) Summary of repression abilities of different dFnCpf1 and dLbCpf1 variants. Repression fold is calculated as the ratio between fluorescence of the positive control and the test systems at 10^3^µM IPTG inducer concentration in (C) and (D). (C) Repression curves of three dLbCpf1 variants. The positive control (“P”) was of the strain with an empty crRNA plasmid, while the negative control (“N”) shows the background fluorescence of a strain with an empty *gfp* plasmid. (D) Repression curve of three dFnCpf1 variants. The positive and negative controls are the same as in (C). Error bars represent standard deviation of fluorescence for three independent experiments on different days. For crRNA sequences see Table S3.

### Minimal crRNA length requirements for dCpf1’s regulatory activity

A unique function of Cpf1 is crRNA processing, where pre-crRNA containing multiple units of a 36nt direct repeat (DR) followed by gRNA is cleaved and truncated to mature crRNAs of a 19nt DR-gRNA structure. In several Class I CRISPR systems, crRNA processing is carried out by an independent Cas6-family of ribonucleases, and this process is required for the subsequence assembling of a functional Cas endonuclease complex on crRNA. To find out if crRNA processing is essential for the gene regulatory function of dCpf1, we expressed crRNAs of various DR lengths ranging from 16nt to 36nt in the reporter system (Figure 2A). Interestingly, we found that all crRNAs with DR length >19nt showed the same repression activity as the crRNA with DR length of exactly 19nt. Since the latter did not undergo processing, we concluded that the regulatory activity of dCpf1 is independent of its crRNA processing activity. We also found that crRNAs with DR length <19nt were not able to direct gene repression (Figures 2B&C). This is in contrast with a previous *in vitro* experiment that showed for Cpf1, crRNA with DR lengths of 16-18nt were still able to induce target DNA cleavage (15).

**Figure 2.**
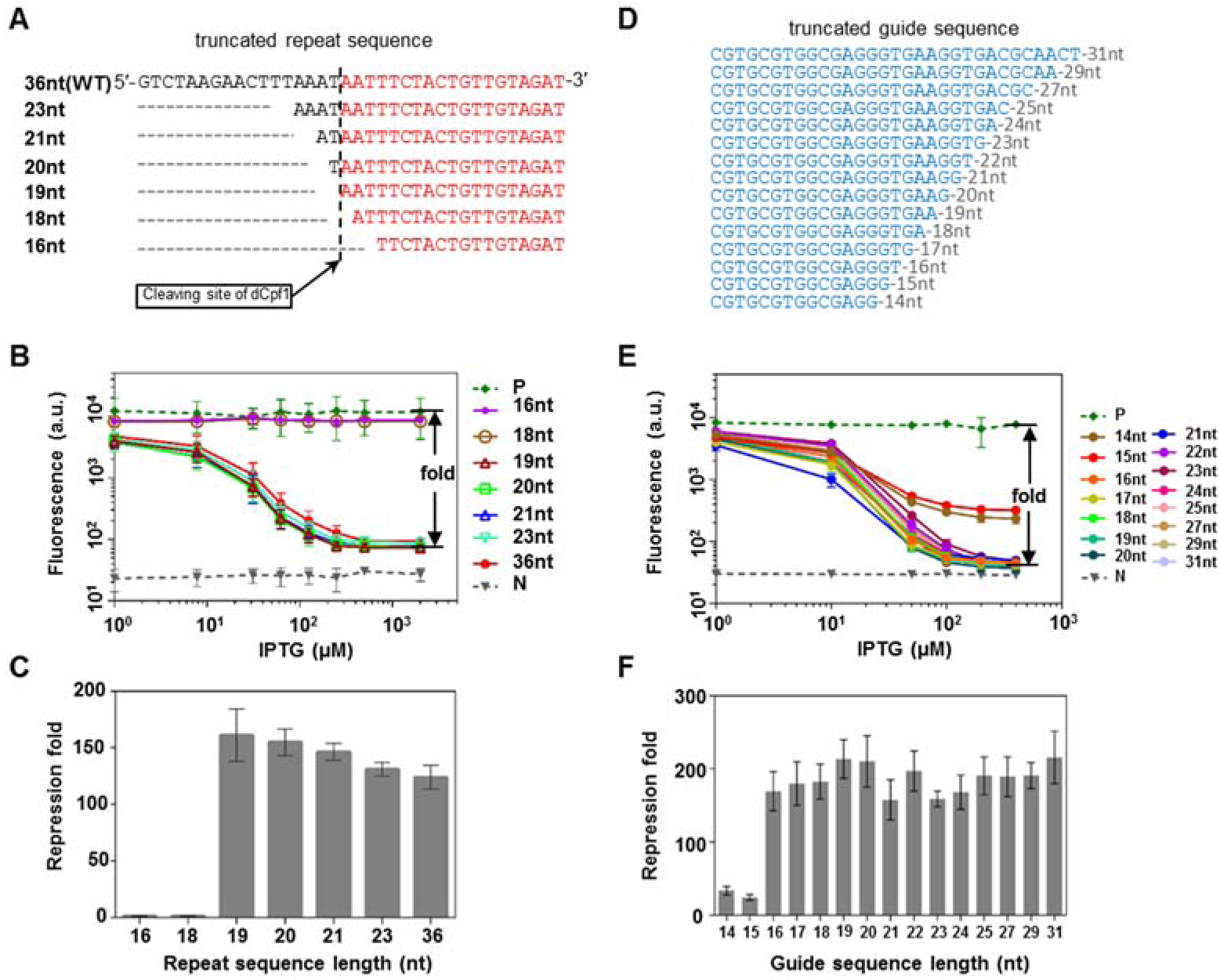
The effect of repeat and guide sequence lengths on gene repression by dFnCpf1-crRNA. (A) Aligned repeat sequences of different lengths used in crRNAs. The dashed line indicates the cleavage site on crRNA during crRNA processing by dFnCpf1. Red colored sequences remain in the mature crRNA, while the rest of the sequences are cleaved off. (B) Gene repression curves of dFnCpf1 with truncated repeat sequences. (C) Maximal repression folds of dFnCpf1 with the same set of truncated repeat sequence as in (B). (D) Aligned guide sequences of different lengths used in crRNAs. (E) The repression curves for different lengths of guide sequences in the dFnCpf1-crRNA system. (F) Maximal repression folds of dFnCpf1 with the same set of truncated guide sequences as in (E). Error bars represent standard deviation of fluorescence for three independent experiments on different days. Positive and negative controls are the same as in Figure l.

Another functional element in crRNA is the guide sequence whose length is believed to be a crucial parameter for the DNA cleaving efficiency of the Cpf1 nuclease. Cpf1 generates mature crRNAs with guide sequence of typically 24nt long. We examined how the extension and truncation of the guide sequence affect the regulatory efficiency of dCpf1 by constructing a number of guide sequences with lengths from 14nt to 31nt (Figure 2D). The results showed a guide sequence ≥ 16nt was able to elicit 200-fold gene repression, while the 14- and 15-nt guide sequences elicited only 20-fold repression (Figure 2E&F). For Cpf1, a previous study suggested a threshold guide sequence length of 18nt below which DNA targeting and cleavage was not observed (15). These results together suggested a 16-18nt minimal guide sequence length required for DNA targeting.

### Enhanced gene repression through multiplex targeting of dCpf1

Next, we sought to demonstrate the ability of multiplex gene regulation of the CRISPR-dCpf1 system. As the targeting of multiple genes has been demonstrated in several recent studies (17,18,20), and a single bound dCpf1, without dedicated inactivation domains, was not sufficient in suppressing gene expression in human HEK293T cells (20), we studied gene repression by tandemly positioned dCpf1 roadblocks within a single gene. Guide sequences were selected to target three independent segments within the coding region of the *sf-gfp* gene. These 24nt guide sequences were then connected by the 36nt DR sequence and co-expressed under a constitutive promoter (Figure 3A). Gene repression is achieved through interference of the bound dCpf1 proteins with the proceeding RNA polymerase complex. We found that crRNAs targeting any one of the three segments resulted in varied but significant gene repression (10~100-fold). Repression was further augmented by doubly or triply combined crRNAs, presumably through a stronger blockage of transcription elongation by the tandemly loaded dCpf1 (Figure 3C). Strikingly, the triply combined crRNAs completely abolished *gfp* expression (>300-fold reduction). The fold reduction by multiplex targeting, relative to individual targetings, was between additive and multiplicative. These results suggested that dCpf1 can process co-transcribed guide sequences targeting multiple DNA segments, conferring combinatorially augmented gene repression.

**Figure 3.**
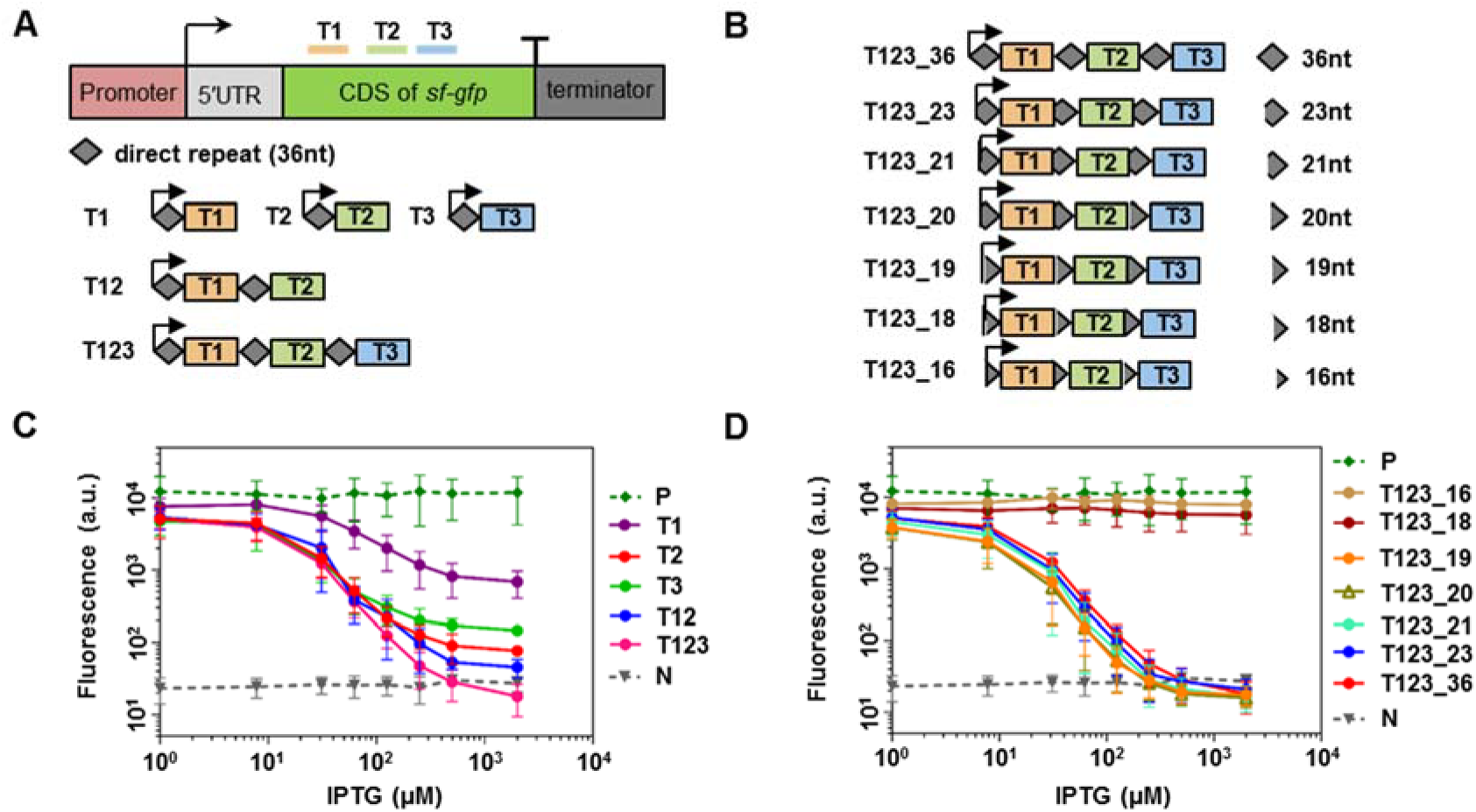
Repression by dFnCpf1 with co-transcribed crRNAs. (A) Schematic representation of target sequences for each single crRNA, as well as the design of individual and combined multiple crRNAs. (B) Different lengths of repeat sequence in the triply-combined crRNA co-transcript. (C) Repression curves of dFnCpf1 with single or multiple crRNAs. (D) Repression curves of dFnCpf1 with varied repeat sequence lengths in the same triply-combined crRNA co-transcript. Error bars represent standard deviation of fluorescence for three independent experiments on different days. Positive and negative controls are the same as in Figure 1. For crRNA sequences see Table S3.

Co-transcription of multiple crRNAs ensures uniform expression among all gRNAs, and reduces the genetic instability associated with the repeated use of promoters and terminators. Yet, in the crRNA coding region, a repeat structure conferred by the DR sequences could also lead to genetic instability through an increased chance of homologous recombination as the length of DR increases (26). We further optimized the system by truncating the interspersed DR sequences, and identified the minimal DR length essential for multiple DNA targeting (Figure 3B). In consistence with the condition for single crRNAs, we found a 19nt DR is required for dCpf1-mediated multiplex repression (Figure 3D).

### dFnCpf1’s regulatory activity strongly depends on the PAM sequence

Previous studies have shown a strong dependence of CRISPR activity on the PAM sequence. For FnCpf1, CTN and TTN were identified as the preferred PAM sequences for DNA cleavage. We selected two sets of targets on both the template and non-template strands of the *sf-gfp* gene based on these motifs, and tested the gene repression activity of dFnCpf1 (Figure S1A). We observed that none of the non-template strand targets generated significant repression (Figure S1B) – a strand bias also reported in other studies – while the template strand targets showed a broad range of repression strengths (Figure S1C). Unlike the case for dCas9 (Figure S2A), for dCpf1, repression strengths were not correlated with the targets’ locations within the coding region (Figure S2B), suggesting factors other than transcript length significantly influenced dCpf1’s regulatory activities. We further selected three sets of targets, each containing three targets starting from a T-rich region, but shifted by 1- or 2-nt relative to each other. Targets selected this way had similar distances from the transcription start site and similar base compositions, and all had TTN as the PAM sequence. However, within each set, repression activities were still drastically different (Figure 4). These results were strongly indicative of TTN as an incomplete characterization of the PAM sequence preference for dFnCpf1, and we speculated that the bases adjacent to the core TTN (and perhaps CTN) motif may underlie the discrepancies in dFnCpf1’s regulatory activity. For example, in Figure 4A, the extended PAMs were G<u>TT</u>T, T<u>TT</u>T, and T<u>TT</u>C, respectively. While the TTTC PAM showed over 100-fold repression, the GTTT PAM was unable to repress gene expression at detectable levels.

**Figure 4.**
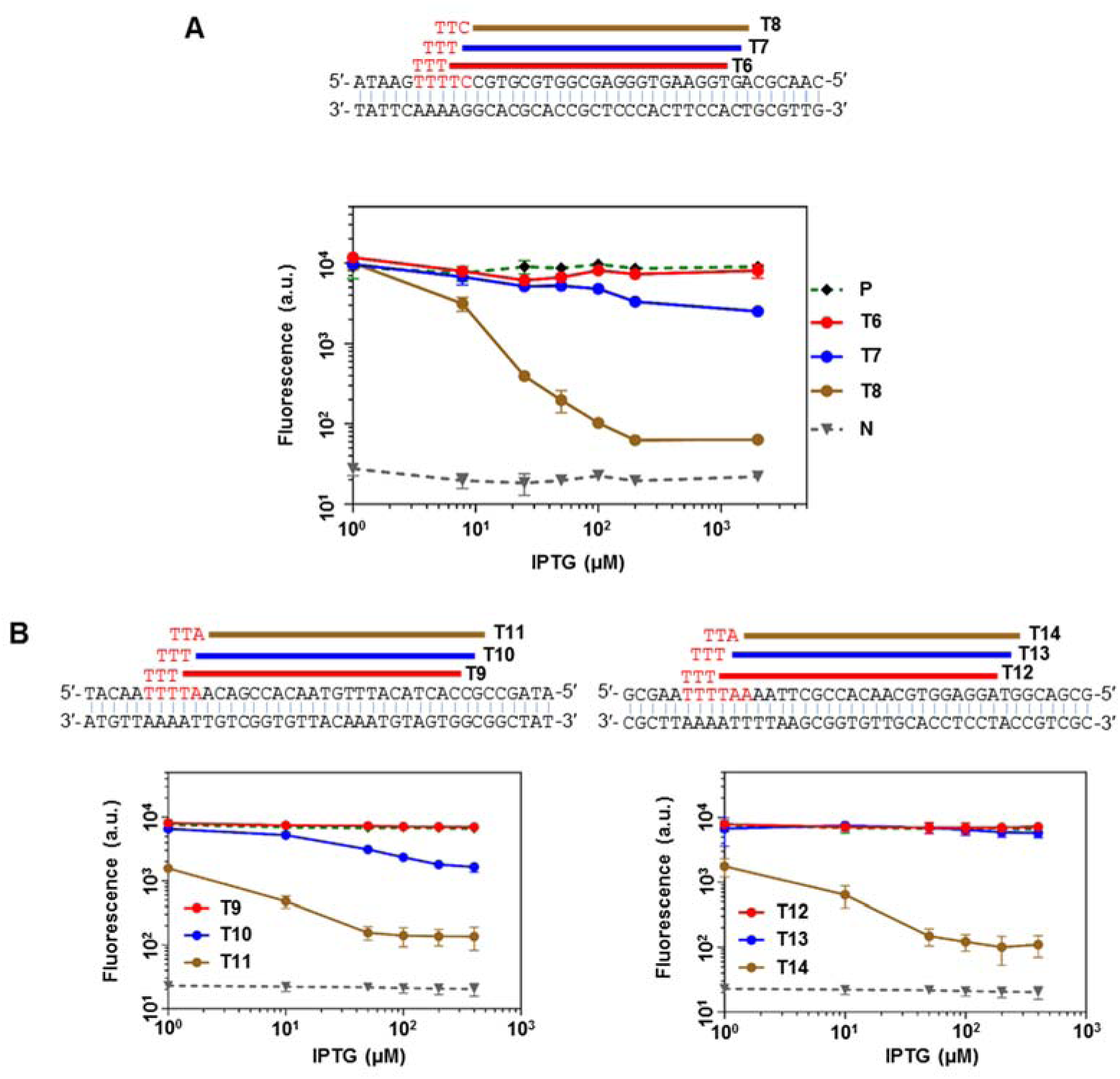
Gene repression on sliding targets with the canonical TTN PAM motif for dFnCpf1. (A) Top panel: targets T6-T8 were selected within the *gfp* coding sequence by 1-nt or 2-nt shifting. Red letters show the corresponding PAM sequences. Lower panel: gene repression by dFnCpf1 targeting the respective sequences. (B) Another two sets of 1-nt or 2-nt shifted target sequences and the respective repression curves by dFnCpf1. Error bars represent the standard deviation of fluorescence for three independent experiments on different days. Positive and negative controls are the same as in Figure 1. For crRNA sequences see Table S3.

### Systematically investigating the effect of PAM sequence for dFnCpf1 and dLbCpf1

To reveal the full range of regulatory activities conveyed by PAM variation, we constructed a library of cells harboring the negative reporter system, with dCpf1 target sequence insertions varying in a randomized 6bp tract as the PAM sequence. The insertion was placed in the 5’-UTR region of the *yfp* gene and followed by a ribozyme-based insulator (27), such that difference in the PAM sequences would not interfere with basal transcription or translation efficiency in the absence of dCpf1 (Figure 5A). Indeed, in Figure 5B, under the non-induced condition, the flow cytometry measured fluorescence distributions of cells harboring the randomized PAM-library (grey line), of the construct with the previously proposed PAM (black dashed line) and of five constructs with mutated PAMs (colored lines) all collapsed onto one curve, indicating the effect of randomized PAM sequences had been successfully eliminated. The library was then subjected to three rounds of dCpf1 induction and fluorescence sorting, from which process, clones showing dramatically varied *yfp* expression levels were randomly picked and sequenced at the PAM locus (Figure S4). Table S1 lists 200 and 133 non-redundant PAM sequences identified in the screens for dFnCpf1 and dLbCpf1, respectively (Figure 5C & Figure S5A). We further measured the fluorescence of these clones at different inducer concentrations (20μM, 50μM and l00µM, IPTG, Figure 5D & S5B). A power law scaling was observed between fluorescence at high and low inducer concentrations when PAM strength was weak or moderate, in consistence with a simple gene expression model depending on dCpf1 concentrations. As PAM became strong, repression levels gradually saturated along both axes.

**Figure 5.**
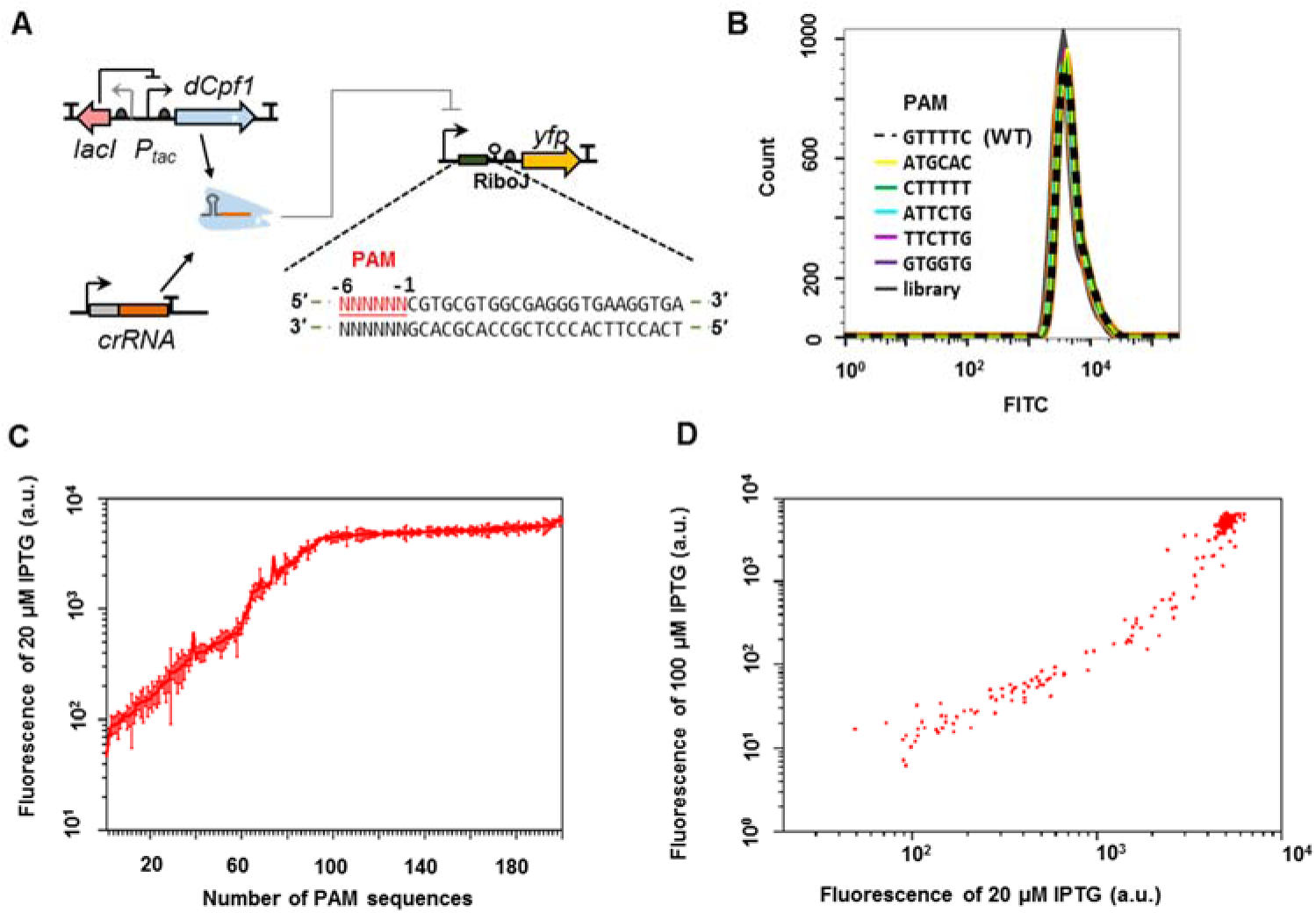
Negative reporter screen for the PAM dependence of dFnCpf1’s regulatory activity. (A) Design of the screening circuit. A randomized 6-nt PAM sequence (red) was placed upstream of a fixed target sequence, and a ribozyme insulator (RiboJ) was inserted between the target and the reporter gene to eliminate the effect of PAM sequences on *yfp* translation. (B) Fluorescence distributions measured by flow cytometry for clones carrying the specified PAM sequences or cell populations carrying the randomized PAM library, under the non-induced condition. (C) Fluorescence measured under the induced condition, for n=200 clones carrying different PAMs randomly selected from flow-cytometer sorted library cells. Error bars represent the standard deviation of two to four independent experiments on different days. (D) Fluorescence for the 200 sample clones under two different IPTG inducer concentrations (20 µM and 100 μM)

These results suggest that for the CRISPR-dCpf1 system, variations in the PAM sequences could produce a large dynamic range for gene expression regulation. In contrast to the irreversible DNA cleavage reaction for which any “good” PAM would suffice, gene regulation applications could take advantage of a more nuanced activity difference between PAMs to achieve controllable outputs. However, as our entire PAM library contained 4,096 sequences, it was both impractical and uneconomical to screen and sequence all clones. Therefore, we designed an interpolation algorithm to predict PAM strengths using information gathered from a small sample pool, such as the 200 dFnCpf1 clones picked by fluorescence levels. The algorithm is based on the assumption of a semi-smooth regulatory strength landscape in the PAM sequence space, in other words, the regulatory strength of a PAM sequence of length *k* is computed as the average strengths of all PAMs that are different by one nucleotide at only one of the *k* locations. Strength information at location *i* (*i*=1…*k*) is weighted by the *degeneracy* of the location. A non-degenerate location is a location where variations in base identity have exhibited very different regulatory outputs in the sample set, and thus all information at this location is discarded for predictive purposes. Whenever possible, context dependency is considered in evaluating location degeneracy. A detailed explanation of the algorithm can be found in Supplementary Information. Unlike the conventional PWM model or sequence logo methods, the algorithm does not assume positional independence between bases, and therefore, it automatically captures all sequence patterns and features contained by the sample pool.

We predicted PAM strengths for all 6nt words based on data from 200 samples for dFnCpf1 and 133 samples for dLbCpf1 (Figure 6A, S6A & Table S2). Conversely, we used the predicted values for unmeasured words to back-predict strengths of measured PAMs. This yielded a >0.99 correlation with measured values, indicating a minimal loss of information through the course of interpolation (data not shown). The results indicated that in general, for both dFnCpf1 and dLbCpf1, positions 1 and 2 did not had significant effects on PAM strengths. For dFnCpf1, PAM strength was most sensitive to the 4- and 5-th location, while position 3 contributed to PAM strength diversity more than position 6. For dLbCpf1, positions 3-6 all affected PAM strength strongly (Figure 6B & S6B). When ranking samples based on repression activity, we found that for dFnCpf1, the strongest PAMs were (TT)TTTV and (T)TTV, whereas T was strongly disfavored at the last position. The other previously identified CTN motif generated only moderate repression activities (Figure 6A). For dLbCpf1, the strongest repressions were elicited by (T) TTTV PAMs, followed by CTTV. Like dFnCpf1, there was a strong preference against T at the last position in strong and moderate PAMs. However, for dLbCpf1, TTTT was able to induce medium repression, with a 5’-T further enhancing its activity (Figure S5A and S6A).

**Figure 6.**
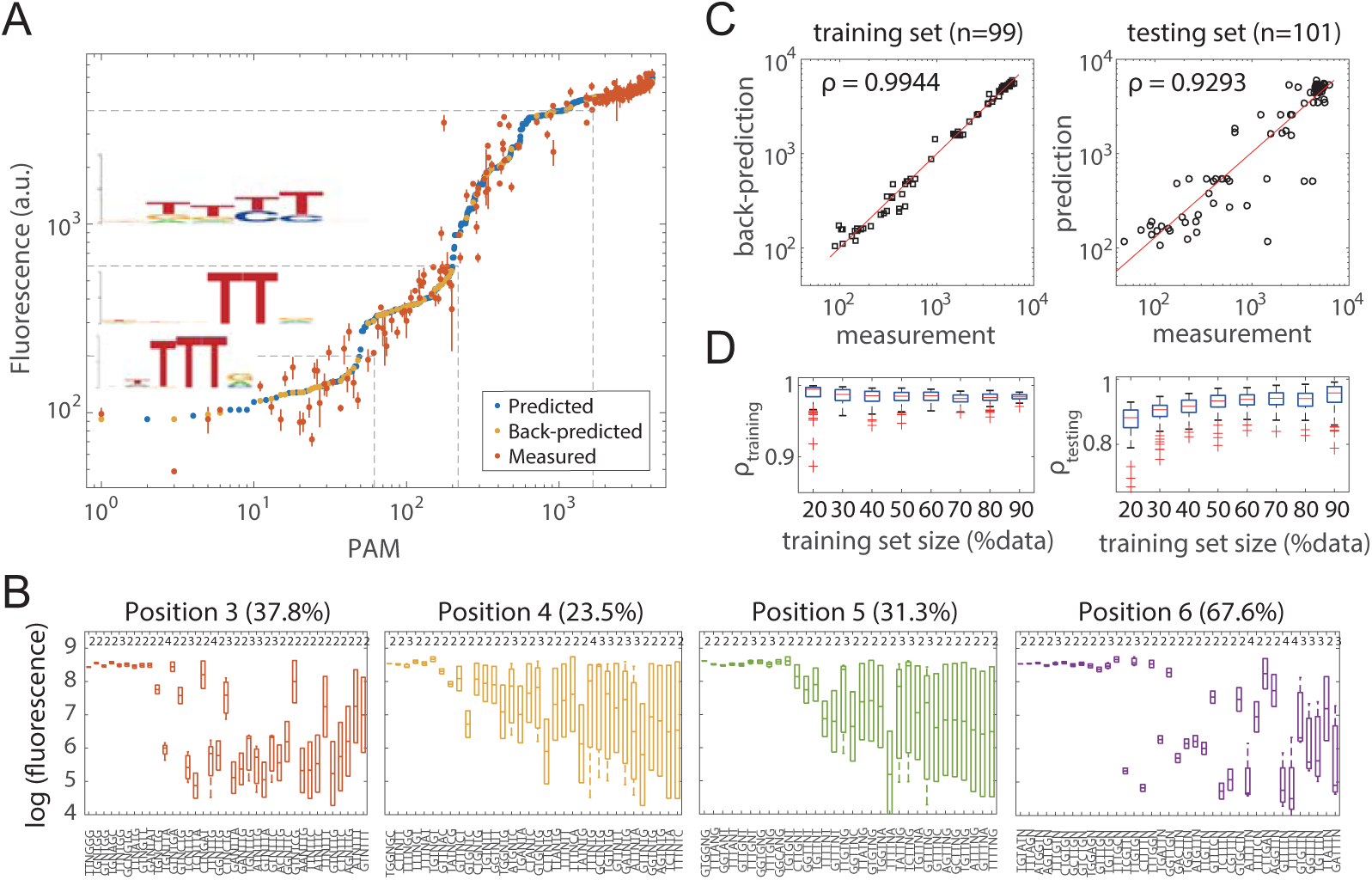
Predictions of PAM strengths for dFnCpf1. (A) Predictions of repression strengths for 4,096 6-nt PAMs. PAMs are sorted by predicted values. Back predictions were made for measured PAMs in the sample pool (n=200) from values predicted for unmeasured PAMs (n=3,896). For measured values (red dots), error bars show the standard error of mean for two to four independent measurements on different days. Sequence logos were obtained from measured PAMs with fluorescence strengths in ranges (0,200), (200, 600) and (600, 4000). (B) Site degeneracy for PAM positions 3-6. Box plots for measured PAM strengths (log fluorescence values, *y*-axis) of the sequence context specified on the x-axis. Percentages in parentheses indicate fractions of contexts that are degenerate by a 2-fold threshold. Numbers on top indicate the number of measured words within each sequence context. (C) Correlation between predictions and measured values in a sample run of cross-validation tests at 50% training set-testing set split. *ρ*: Pearson correlation between log values. (D) Summary of cross-validation tests. Correlation between predictions and measurements for the training (left) and testing (right) sets. In each run, data for training were randomly selected from and thus have the same distribution as the sample pool. 100 runs were conducted for each training set-testing set splitting ratio.

To evaluate the predictive power of our algorithm, cross-validation was done by splitting the sample pool for dFnCpf1 into training and testing sets, at proportions from 20% to 90% (for the training set). When using 50% randomly selected samples as the training set, the predictions for the testing set were >0.90 correlated with measured values, and back-prediction showed >0.95 correlations with data in the training sets (Figure 6C). Even with only 20% (n=40) of the sampled PAMs as the training data, a correlation >0.8 could be obtained with the testing set (Figure 6D). These numbers decreased mildly for dLbCpf1, whose sample pool were smaller and less biased toward high and low repression ranges (Figure S6C & S6D). When we applied a strictly uniform selection method in the low, medium, and high repression ranges, irrespective of the repression strength distribution of the original sample pools, correlation between predictions and the testing sets were around 0.75 (at 53% data as training set) for dFnCpf1 and 0.4 (at 14% data as training set) for dLbCpf1 (Figure S7). These results underscored the importance of PAM sequences sampled at high and low repression ranges, which should generate sufficient information for the algorithm to successfully interpolate for any other PAM sequence.

## Discussion

In this article, we systematically investigated the key constraints and properties of the CRISPR-dCpf1 system as transcriptional repressors in *E. coli* cells. In comparison to the dCas9 based CRISPR systems, dCpf1 offers the unique potential of multiplex gene regulation with its ability to autonomously process crRNA co-transcripts and subsequently target multiple independent DNA sequences. This ability minimizes the uncertainty in crRNA relative dosages and genetic stabilities, as previously seen in systems with dCas9 and independently transcribed crRNAs. This is key to large scale standardized perturbation experiments such as whole transcription network engineering. There have recently been multiple reports on dCpf1’s gene regulation applications in bacteria, plants, and human cells. Although repression in bacteria was attained, repression in *Arabidopsis* and activation in human HEK293T cells was unstable and idiosyncratic. A systematic characterization of the CRISPR-dCpf1 system with respect to its DNA binding properties is obviously in need to further enhance performance in these experimental systems.

We compared the repression activities of dCpf1 mutant forms including single and double mutations at the two previously identified catalytic residues for Cpf1’s DNase activity. For both dFnCpf1 and dLbCpf1, double mutations compromised regulatory activities. Between the two single mutation variants, D917A/D832A generated consistently strong regulatory activity, whereas E1006 in dFnCpf1 was much less efficient in DNA binding than D917A. This suggests that the RuvC-I domain mutation generally do not interfere with DNA binding, whereas the mutation in RuvC-II may in some dCpf1 proteins affect dCpf1-crRNA-DNA complex formation.

We found that for dCpf1, crRNA cleavage was not essential for subsequent DNA targeting. The wild type crRNAs adopt a 19nt DR-24nt gRNA form. In our studies, the minimal length requirements were 19nt for the direct repeat and 16nt for the guide sequence. In a previous *in vitro* DNA cleavage assay, 16-18nt direct repeat sequences were able to induce detectable cleavage (15). A previous biochemical study revealed the importance of a 5’-AAU-3’ sequence at the -19 location of the processed crRNA (14). This tri-base region may help stabilizing the dCpf1-crRNA complex. With shorter crRNAs, Cpf1 may still form transient complexes with DNA and produce strand breaks *in vitro*. However, tight binding of dCpf1-crRNA to the DNA target demanded an intact 19nt direct repeat sequence according to our results.

We further demonstrated a way of enhancing gene repression by using co-transcribed crRNAs targeting DNA sequences located in tandem in the coding region. Multiplex targeting elicited almost complete repression compared to individual targets. Again, a ≥19nt DR length is required in the crRNA co-transcript for crRNA processing and the targeting to the respective sequences.

We found the PAM sequence to be a major factor determining gene repression activity. The previously identified TTN and CTN motifs for FnCpf1 in DNA cleavage assays did not explained the PAM preference in terms of gene regulation by dFnCpf1. Although a T-rich PAM for Cpf1 greatly expands the genomic regions that could be targeted for cleavage, gene regulatory response was sensitively dependent on the exact PAM sequences used. On the same target sequence, a wide range of repression folds were observed when different 6nt preceding sequences were used. We designed a negative reporter screen to identify PAM sequences eliciting strong, medium and weak repressions. We further developed an interpolation algorithm based on context-dependent sequence similarities, using which, we predicted regulatory strengths for all 6nt sequences as PAMs based on measurements of 200 and 133 PAMs for dFnCpf1 and dLbCpf1, respectively. Our analyses suggested for both enzymes a general 4nt core sequence dependence, with T strongly disfavored in the last position, and slightly favored at the proceeding 2nt positions. Specifically, dFnCpf1 and dLbCpf1 both displayed a preference for TTTV PAMs; while for dLbCpf1, other PAMs also emerged as mediating strong regulatory responses. TTTV was previously identified for LbCpf1 (22), and recently identified in a study on genome editing by FnCpf1 in Baker’s yeast (28), while we were preparing this manuscript. This suggests that the difference in strengths of extended PAMs may also be relevant when cleavage is concerned, especially for improving CRISPR DNases that did not function well in certain systems. In (28), the authors claimed that targets with TTTA and (CT) TTTC PAMs did not lead to genome editing, despite conforming to the TTTV motif. Our data suggest a range of 50-300 fluorescence for NNTTTA PAMs and a ~110 fluorescence for CTTTTC. Although these are strong repressions in the 6nt library, the six-fold difference might still significantly affect reaction outcome.

For multiplex gene regulation applications, these predicted PAM sequence strengths for dCpf1 enable the design and implementation of differential regulatory responses among targets at a single dCpf1 induction level. Unlike modulation by inducer concentrations, independent modulation by PAM strengths grants much flexibility for quantitative assessment of complex transcription networks. Moreover, the screening method we developed could be utilized to introduce a control element in arbitrary genes. Compared to targets in the upstream promoter regions, insertions within the 5’UTR region followed by an insulator could minimize the interference on background gene expression level. Compared to targets in the coding sequences, PAM sequences and target sequences in the inserted fragment can be designed separately to achieve desired repression outputs with high specificity. Besides dFnCpf1 and dLbCpf1, dCpf1s from *Acidaminococcus sp*. and *Eubacterium eligens* have also been tested in bacterial and eukaryotic cells (17-20). Our screening and prediction methods could serve a pipeline facilitating the transformation of these proteins into powerful tools for diverse application of multiplex gene regulation.

## SUPPLEMENTARY DATA

Supplementary Data are available at

### ACKNOWLEDGEMENTS

We thank Lili Ji (Peking University), Tingting Li (Peking University) and Junying Jia (Institute of Biophysics, CAS) for technical assistance of flow cytometry.

## References

1. Cho, S.W., Kim, S., Kim, J.M. and Kim, J.S. (2013) Targeted genome engineering in human cells with the Cas9 RNA-guided endonuclease. Nature biotechnology, 31, 230–232.

2. Cong, L., Ran, F.A., Cox, D., Lin, S., Barretto, R., Habib, N., Hsu, P.D., Wu, X., Jiang, W., Marraffini, L.A. et al. (2013) Multiplex genome engineering using CRISPR/Cas systems. Science, 339, 819–823.

3. Jiang, W., Zhao, X., Gabrieli, T., Lou, C., Ebenstein, Y. and Zhu, T.F. (2015) Cas9-Assisted Targeting of CHromosome segments CATCH enables one-step targeted cloning of large gene clusters. Nat Commun, 6, 8101.

4. Jinek, M., Chylinski, K., Fonfara, I., Hauer, M., Doudna, J.A. and Charpentier, E. (2012) A programmable dual-RNA-guided DNA endonuclease in adaptive bacterial immunity. Science, 337, 816–821.

5. Qi, L.S., Larson, M.H., Gilbert, L.A., Doudna, J.A., Weissman, J.S., Arkin, A.P. and Lim, W.A. (2013) Repurposing CRISPR as an RNA-guided platform for sequence-specific control of gene expression. Cell, 152, 1173–1183.

6. Didovyk, A., Borek, B., Hasty, J. and Tsimring, L. (2016) Orthogonal Modular Gene Repression in Escherichia coli Using Engineered CRISPR/Cas9. Acs Synth Biol, 5, 81–88.

7. Nielsen, A.A. and Voigt, C.A. (2014) Multi-input CRISPR/Cas genetic circuits that interface host regulatory networks. Molecular systems biology, 10, 763.

8. Cress, B.F., Toparlak, O.D., Guleria, S., Lebovich, M., Stieglitz, J.T., Englaender, J.A., Jones, J.A., Linhardt, R.J. and Koffas, M.A. (2015) CRISPathBrick: Modular Combinatorial Assembly of Type II-A CRISPR Arrays for dCas9-Mediated Multiplex Transcriptional Repression in E. coli. Acs Synth Biol, 4, 987–1000.

9. Lv, L., Ren, Y.L., Chen, J.C., Wu, Q. and Chen, G.Q. (2015) Application of CRISPRi for prokaryotic metabolic engineering involving multiple genes, a case study: Controllable P(3HB-co-4HB) biosynthesis. Metabolic engineering, 29, 160–168.

10. Elhadi, D., Lv, L., Jiang, X.R., Wu, H. and Chen, G.Q. (2016) CRISPRi engineering E. coli for morphology diversification. Metabolic engineering, 38, 358–369.

11. Li, S., Jendresen, C.B., Grunberger, A., Ronda, C., Jensen, S.I., Noack, S. and Nielsen, A.T. (2016) Enhanced protein and biochemical production using CRISPRi-based growth switches. Metabolic engineering, 38, 274–284.

12. Zalatan, J.G., Lee, M.E., Almeida, R., Gilbert, L.A., Whitehead, E.H., La Russa, M., Tsai, J.C., Weissman, J.S., Dueber, J.E., Qi, L.S. et al. (2015) Engineering complex synthetic transcriptional programs with CRISPR RNA scaffolds. Cell, 160, 339–350.

13. Nissim, L., Perli, S.D., Fridkin, A., Perez-Pinera, P. and Lu, T.K. (2014) Multiplexed and programmable regulation of gene networks with an integrated RNA and CRISPR/Cas toolkit in human cells. Mol Cell, 54, 698–710.

14. Fonfara, I., Richter, H., Bratovic, M., Le Rhun, A. and Charpentier, E. (2016) The CRISPR-associated DNA-cleaving enzyme Cpf1 also processes precursor CRISPR RNA. Nature, 532, 517–521.

15. Zetsche, B., Gootenberg, J.S., Abudayyeh, O.O., Slaymaker, I.M., Makarova, K.S., Essletzbichler, P., Volz, S.E., Joung, J., van der Oost, J., Regev, A. et al. (2015) Cpf1 is a single RNA-guided endonuclease of a class 2 CRISPR-Cas system. Cell, 163, 759–771.

16. Zetsche, B., Heidenreich, M., Mohanraju, P., Fedorova, I., Kneppers, J., DeGennaro, E.M., Winblad, N., Choudhury, S.R., Abudayyeh, O.O., Gootenberg, J.S. et al. (2017) Multiplex gene editing by CRISPR-Cpf1 using a single crRNA array. Nature biotechnology, 35, 31–34.

17. Kim, S.K., Kim, H., Ahn, W.C., Park, K.H., Woo, E.J., Lee, D.H. and Lee, S.G. (2017) Efficient Transcriptional Gene Repression by Type V-A CRISPR-Cpf1 from Eubacterium eligens. Acs Synth Biol.

18. Tak, Y.E., Kleinstiver, B.P., Nunez, J.K., Hsu, J.Y., Horng, J.E., Gong, J., Weissman, J.S. and Joung, J.K. (2017) Inducible and multiplex gene regulation using CRISPR-Cpf1-based transcription factors. Nat Methods, 14, 1163–1166.

19. Tang, X., Lowder, L.G., Zhang, T., Malzahn, A.A., Zheng, X., Voytas, D.F., Zhong, Z., Chen, Y., Ren, Q., Li, Q. et al. (2017) A CRISPR-Cpf1 system for efficient genome editing and transcriptional repression in plants. Nat Plants, 3, 17018.

20. Zhang, X., Wang, J., Cheng, Q., Zheng, X., Zhao, G. and Wang, J. (2017) Multiplex gene regulation by CRISPR-ddCpf1. Cell Discov, 3, 17018.

21. Yamano, T., Nishimasu, H., Zetsche, B., Hirano, H., Slaymaker, I.M., Li, Y., Fedorova, I., Nakane, T., Makarova, K.S., Koonin, E.V. et al. (2016) Crystal Structure of Cpf1 in Complex with Guide RNA and Target DNA. Cell, 165, 949–962.

22. Kim, H.K., Song, M., Lee, J., Menon, A.V., Jung, S., Kang, Y.M., Choi, J.W., Woo, E., Koh, H.C., Nam, J.W. et al. (2017) In vivo high-throughput profiling of CRISPR-Cpf1 activity. Nat Methods, 14, 153–159.

23. Leenay, R.T. and Beisel, C.L. (2017) Deciphering, Communicating, and Engineering the CRISPR PAM. J Mol Biol, 429, 177–191.

24. Leenay, R.T., Maksimchuk, K.R., Slotkowski, R.A., Agrawal, R.N., Gomaa, A.A., Briner, A.E., Barrangou, R. and Beisel, C.L. (2016) Identifying and Visualizing Functional PAM Diversity across CRISPR-Cas Systems. Mol Cell, 62, 137–147.

25. Watkins-Chow, D.E., Varshney, G.K., Garrett, L.J., Chen, Z., Jimenez, E.A., Rivas, C., Bishop, K.S., Sood, R., Harper, U.L., Pavan, W.J. et al. (2017) Highly Efficient Cpf1-Mediated Gene Targeting in Mice Following High Concentration Pronuclear Injection. G3 (Bethesda), 7, 719–722.

26. Chen, Y.J., Liu, P., Nielsen, A.A., Brophy, J.A., Clancy, K., Peterson, T. and Voigt, C.A. (2013) Characterization of 582 natural and synthetic terminators and quantification of their design constraints. Nat Methods, 10, 659–664.

27. Lou, C., Stanton, B., Chen, Y.J., Munsky, B. and Voigt, C.A. (2012) Ribozyme-based insulator parts buffer synthetic circuits from genetic context. Nature biotechnology, 30, 1137–1142.

28. Swiat, M.A., Dashko, S., den Ridder, M., Wijsman, M., van der Oost, J., Daran, J.M. and Daran-Lapujade, P. (2017) FnCpf1: a novel and efficient genome editing tool for Saccharomyces cerevisiae. Nucleic acids research, 45, 12585–12598.

